# TRPV1+ neurons promote cutaneous immunity against *Schistosoma mansoni*

**DOI:** 10.1101/2025.02.06.636930

**Authors:** Juan M. Inclan-Rico, Adriana Stephenson, Camila M. Napuri, Heather L. Rossi, Li-Yin Hung, Christopher F. Pastore, Wenqin Luo, De’Broski R. Herbert

## Abstract

Immunity against skin-invasive pathogens requires mechanisms that rapidly detect, repel or immobilize the infectious agent. While bacteria often cause painful cutaneous reactions, host skin invasion by the human parasitic helminth *Schistosoma mansoni* often goes unnoticed. This study investigated the role of pain-sensing skin afferents that express the ion channel Transient Receptor Potential Vanilloid 1 (TRPV1) in the detection and initiation of skin immunity against *S. mansoni*. Data show that mice infected with *S. mansoni* have reduced behavioral responses to painful stimuli and sensory neurons exposed from infected mice have significantly less calcium influx and neuropeptide release in response to the TRPV1 agonist capsaicin. Using both gain- and loss-of-function approaches, data show that TRPV1+ neurons are critical regulators of *S. mansoni* survival during migration from the skin into the pulmonary tract. Moreover, TRPV1+ neurons were both necessary and sufficient to promote proliferation and cytokine production from dermal γδ T cells as well as neutrophil and monocyte skin accumulation post-infection. These results suggest a model in which *S. mansoni* may have evolved to inhibit TRPV1+ neuron activation as a countermeasure that limits IL-17-mediated inflammation, facilitating systemic dissemination and chronic parasitism.

**One sentence summary:** The parasitic helminth *Schistosoma mansoni* averts IL-17-dependent protective immunity by suppressing skin-innervating TRPV1+ neurons.

## Introduction

The skin is populated by hematopoietic and non-hematopoietic cells that coordinately assemble the first line of defense against pathogen invasion. Certain bacterial and fungal skin infections evoke painful and itchy (pruritic) reactions transmitted by diverse subsets of sensory neurons (afferents) whose cell bodies emanate from dorsal root ganglia (DRG) or trigeminal ganglia (TG)^1–3^. Notably, the predominant subset of pain-sensing neurons (nociceptors) that express the non-selective cation channel transient receptor potential vanilloid member 1 (TRPV1) have been recognized as central regulators of barrier tissue immune responses during infection, autoimmunity, allergy, cancer, wound healing, hair growth, and tolerance^1–13^. TRPV1+ afferents not only respond to canonically stimuli, heat or capsaicin, but can also directly respond to bacterial and/or fungal products. Activation of these neurons leads to secretion of neuropeptides such as calcitonin gene-related peptide (CGRP) and Substance P (SP) that in turn, direct effector functions of skin-resident hematopoietic cells^1,3,14^. While several bacterial species induce TRPV1 neuron-derived CGRP to suppress neutrophil responses to favor bacterial colonization, fungal antigen-induced CGRP from TRPV1+ neurons stimulates myeloid antigen-presenting cells (APC) to induce host-protective IL-17 responses^1–4^. TRPV1+ neurons can also promote detrimental skin inflammation in the context of psoriasis or allergic disease^5,6,12,14^. These studies highlight the importance and context-dependent contributions of sensory neurons in regulating skin immunity.

The blood fluke *Schistosoma mansoni* causes chronic parasitism in ∼250 million people worldwide^15,16^. *S. mansoni* infectious stage larvae (cercariae) are rapidly attracted to mammalian skin and penetrate the host within minutes through the secretion of proteolytic enzymes that degrade extracellular matrix proteins to facilitate their rapid migration into the lung^17,18^. Surprisingly, the cutaneous stage of *S. mansoni* pathogenesis is usually asymptomatic, sometimes causing mild dermatitis and/or itch^15,16,19^. Conversely, zoonotic exposure to other avian schistosome species is marked by severe dermatitis and pruritus, known as Swimmer’s itch, with worsening manifestations upon repeated exposure proposed to prevent systemic parasite dissemination^20–22^. Our recent work demonstrated that itch-transmitting neurons bearing the receptor Mas-related G-coupled protein receptor A3 (MrgprA3) can directly react to *S. mansoni,* but this interaction results in their inactivation^23^. Our study shows that MrgprA3+ neurons can repel *S. mansoni* skin entry and migration through the induction of pro-inflammatory cytokine secretion by myeloid APC that induce IL-17-mediated inflammation. TRPV1 is broadly expressed in DRG and TG nociceptors, with a low-level expression in MrgprA3+ neurons^24,25^, suggesting that the broad population of TRPV1+ neurons may elicit immunity against *S. mansoni*. However, whether nociceptive TRPV1+ neurons contribute to host-protective immune responses against *S. mansoni* has not been addressed.

This work demonstrates that *Schistosoma mansoni* infection reduces thermal pain sensitivity as well as calcium influx and neuropeptide release evoked by TRPV1+ neurons. We postulated that *S. mansoni* may have evolved to suppress TRPV1+ neuron activation that would otherwise elicit host protective skin inflammation thereby favoring parasite invasion and survival. Indeed, local optogenetic TRPV1+ neuron activation prior to *S. mansoni* exposure conferred resistance to skin invasion and attenuated parasite dissemination into the lungs. TRPV1+ neuron activation induced IL-17+, IL-13+, and Ki67+ γδ T cells in skin, increased swelling, and promoted the accumulation of inducible nitric oxide synthetase (iNOS+) neutrophils and monocytes within 1 day post-infection. Conversely, chemical denervation of TRPV1+ neurons increased larval burden in the lungs and diminished γδ T cell, neutrophil, and monocyte responses. Taken together, these data support a model wherein *S. mansoni* suppresses nociceptor effector function(s) that would otherwise initiate host-protective skin inflammation to limit parasite dissemination and chronic parasitism.

## Results

### *S. mansoni* infection disrupts thermal pain sensation and TRPV1 neuron responses

Our recent work indicated that itch-inducing MrgprA3 neurons could directly recognize *S. mansoni* cercarial extracts and drive parasite erradication^23^. Thus, we tested whether *S. mansoni* infection also blocked the functions of the larger subset of the broad population of TRPV1+ sensory afferents. Wild-type (WT) C57BL/6 mice were percutaneously infected in the right (ipsilateral) hindpaw, and thermal pain sensitivity, which is mainly mediated by TRPV1+ nociceptors^26^ was evaluated with the Hargreaves assay 24hrs post-infection. Infrarred-induced withdrawal latency was evaluated in the exposed (ipsilateral) and non-exposed opposite (contralateral) paws and compared to basal conditions in naïve controls (**Fig. 1a**). Data show that *S. mansoni-*infected mice had increased withdrawal times in the ipsilateral (exposed) right paw compared to naïve controls. However, withdrawal latency of contralateral (unexposed) paws of *S. mansoni-*infected mice was not significantly different from naïve controls (**Fig. 1b**). To test whether nociception transmitted by TRPV1+ neurons is reduced following *S. mansoni* infection, primary neuron cultures derived from lumbar DRG that innervate both the ipsilateral and contralateral paws of naïve and *S. mansoni*-infected mice were cultured for 2 days followed by evaluation of calcium influx and neuropeptide release evoked by the TRPV1 canonical ligand capsaicin^27^. As expected, capsaicin evoked robust calcium mobilization in 68% (46/68) of DRG neurons from naïve control mice. However, only 26% (9/34) of DRG neurons isolated from *S. mansoni*-infected mice responded to capsaicin, as shown by reduced area under the curve (AUC) values of capsaicin-induced calcium traces (**Fig. 1c-f**). Consistently, capsaicin exposure induced robust CGRP and SP secretion from DRG neurons of naive mice, but DRG neurons isolated from *S. mansoni*-infected mice had significantly reduced neuropeptide secretion (**Fig. 1g,h**). Conversely, DRG neurons isolated from contralateral (unexposed) side of *S. mansoni*-infected mice did not exhibit altered neuropeptide secretion, suggesting local TRPV1 neuron inhibition. TRPV1 immunofluorescence (IFA) staining was detected in DRG neurons from naïve and *S. mansoni*-infected mice that also co-stained for the ion channel Nav1.8, indicating that reduced capsaicin responsiveness was not associated with reduced TRPV1 expression after *S. mansoni* infection (**Supp. Fig. 1a,b**)^3^. Interestingly, naïve DRG neurons that were exposed to *S. mansoni* cercarial extract (SCE) *in vitro* also released significantly less CGRP in response to capsaicin (**Supp. Fig. 1c**), which is consistent with our published results showing reduction of capsaicin-induced calcium influx on naïve DRG neurons incubated with SCE^23^. These results suggest that exposure to parasitic components during *S. mansoni* infection suppresses the responsiveness of local TRPV1+ neurons.

**Figure 1.**
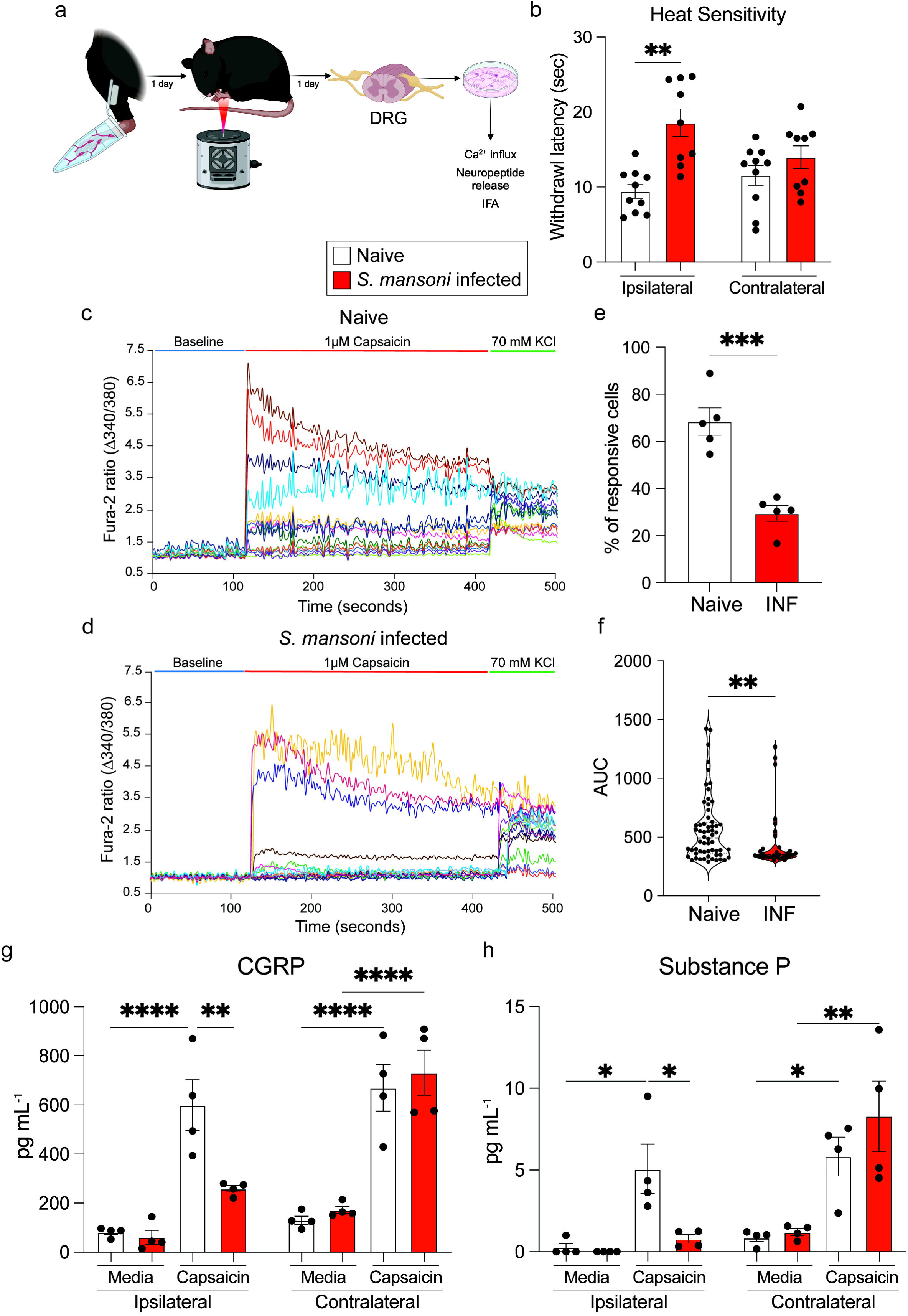
*Schistosoma mansoni* infection reduces TRPV1 neuron responses. (**a**), Experimental approach for percutaneous exposure to *S. mansoni* in the paw followed by assessment of thermal pain sensitivity 1 day after infection and DRG collection 2 days post-infection. DRG cultures were subsequently evaluated for capsaicin-induced calcium influx and neuropeptide release. (**b**), Thermal pain sensitivity in the ipsilateral (exposed) and contralateral (unexposed) paws of naïve and *S. mansoni* infected mice assessed by withdrawal times (seconds) to infrared heat. (**c,d**), Representative image of calcium traces of DRG neuron cultures generated from wildtype naïve or 2-day *S. mansoni*-infected mice. DRG neurons were sequentially treated with 1μM capsaicin for 5 minutes followed by stimulation with 70mM KCl for 1 minute. (**e**), Percentage of capsaicin-responsive DRG neurons per coverslip in naïve and *S. mansoni*-infected mice. (**f**), Area under the curve (AUC) values of intracellular calcium levels of wildtype DRG neurons from naïve or *S. mansoni*-infected mice during capsaicin treatment. (**g,h**), CGRP and Substance P supernatant levels of wildtype DRG neurons generated from wildtype naïve or 2-day *S. mansoni*-infected ipsilateral and contralateral paws treated with vehicle or 1μM capsaicin for 1 hour. *P* values were determined by two-tailed Student’s t-tests, or One-way ANOVA with post hoc correction. *P<0.05, **P<0.01, ***P<0.001, ****P<0.0001. Representative of 2-3 independent experiments, each with ≥4 biological replicates.

### TRPV1+ neuron stimulation protects against *S. mansoni* infection and promotes γδ T cell responses

Given that *S. mansoni* infection disrupted TRPV1+ neuron activation, we postulated that *S. mansoni* inactivates pain-sensing TRPV1+ neurons that would otherwise promote host-protective neurogenic inflammation. To test this hypothesis, a gain-of-function approach was employed to pre-emptively activate TRPV1+ neurons *in vivo* using optogenetics in transgenic mice that expressed channelrhodopsin 2 (ChR2), a light-sensitive non-selective cation channel that evokes action potentials upon exposure to 473 nm blue light^4,28,29^. In ChR2 ^flox/flox^ mice, the *Chr2* gene inserted in the *Rosa26* locus is preceded by a loxP-STOP-loxP cassette. Upon interbreeding with TRPV1^Cre^ mice^30^, Cre expression driven in TRPV1+ neurons removes the STOP cassette, allowing for ChR2 expression^31^. TRPV1^Cre^ ChR2^flox/flox^ (TRPV1-ChR2) or ChR2 ^flox/flox^ littermate control mice were exposed to 473 nm blue light in the ear skin for 5 consecutive days (30 mins/day), followed by percutaneous infection with *S. mansoni* cercariae for 30 min. Skin penetration efficiency and the number of parasites that disseminated to the pulmonary tract were evaluated at 6 days post-infection (**Fig. 2a**). Consistent with previous reports^4^, light exposure in TRPV1-ChR2 mice resulted in increased ear thickness compared to littermate controls (**Fig. 2b**). Light-stimulated TRPV1-ChR2 mice had significantly increased numbers of non-penetrating cercariae and markedly reduced numbers of lung larvae at day 6 when compared to littermate controls (**Fig. 2c,d**). Next, we evaluated the cytokine and activation profile of skin lymphocyte populations by flow cytometry in control or TRPV1-ChR2 mice 1 day after *S. mansoni* infection (**Supp. Fig. 2a**). Control and TRPV1-ChR2 mice presented with similar proportions and numbers of γδ T cells, dendritic epidermal T cells (DETC), CD4+ helper T cells, CD8+ cytotoxic T cells, and group 2 innate lymphoid cells (ILC2s) (**Supp. Fig. 2b**). However, IL-17+, IL-13+, and Ki67+ γδ T cells were increased in TRPV1-ChR2 mice compared to littermate controls following *S. mansoni* infection (**Fig. 2e,f**). Conversely, IL-17, IL-13, and Ki-67 expression were equivalent in CD4+ T helper cells from control and TRPV1-ChR2 mice (**Supp. Fig. 2c-e**). This suggests that preemptive activation of TRPV1 neurons promoted skin γδ T cell responses that prevented *S. mansoni* skin entry and dissemination.

**Figure 2.**
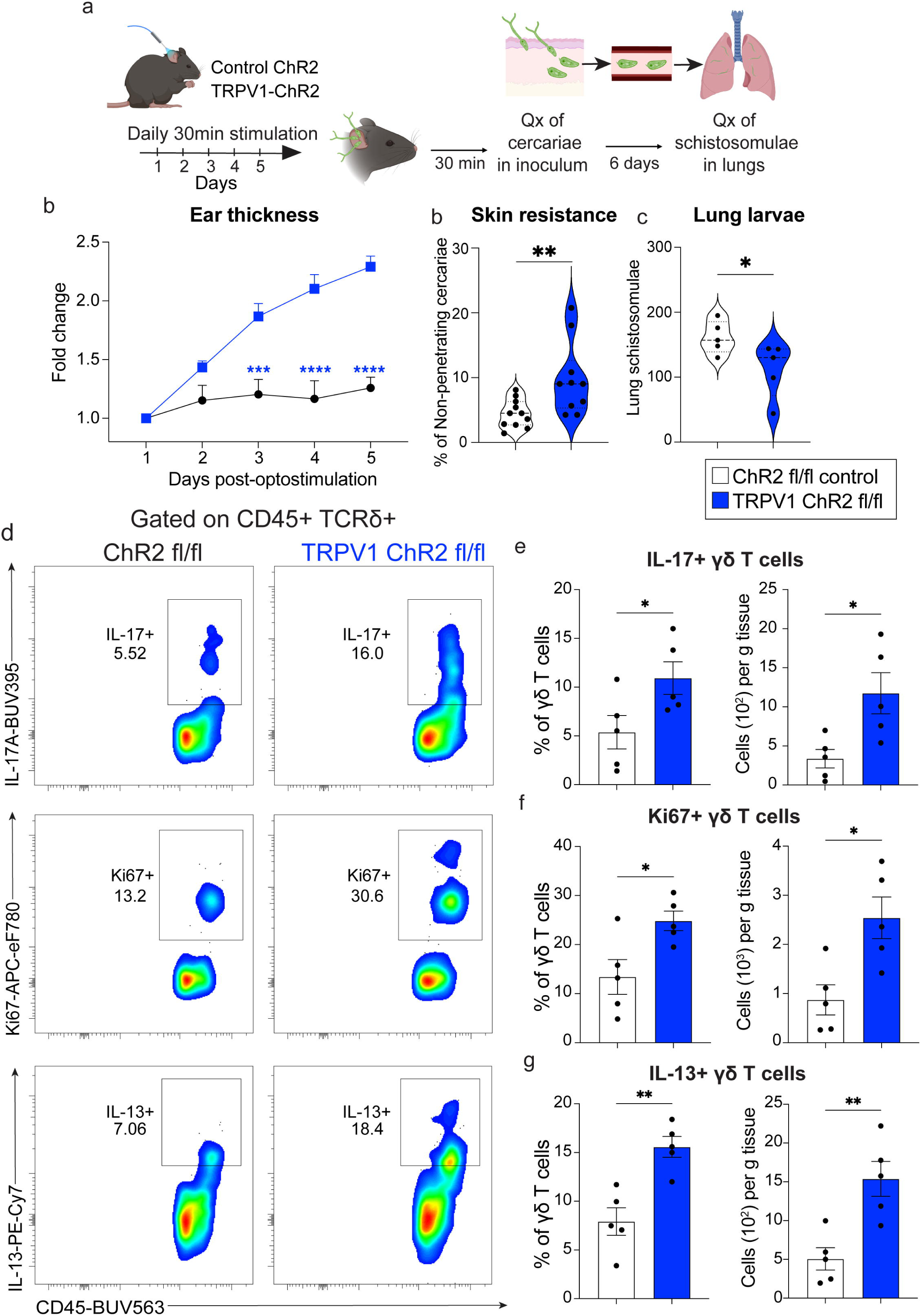
TRPV1 neuron activation confers resistance to *S. mansoni* and induces cytokine expression in γδ T cells. (**a**), Experimental approach for optogenetic ear stimulation of ChR2 control or TRPV1-ChR2 mice followed by evaluation of skin resistance after exposure to 150-200 *S. mansoni* cercariae for 30mins or lung larval (schistosomulae) load 6 days post-infection with 500 *S. mansoni* cercariae. Life cycle is depicted for reference. (**b**), Ear thickness measured daily during photostimulation of ChR2 control or TRPV1-ChR2 mice. (**c**), Percentage of non-penetrating cercariae from ChR2 control or TRPV1-ChR2 mice that were previously photostimulated as shown in A. (**d**), Lung schistosomulae of light-stimulated ChR2 control or TRPV1-ChR2 mice quantified 6 days post-infection. (**e**), Representative dot plots of IL-17, IL-13, and Ki-67 expression in dermal γδ T cells (CD45+ CD90+ TCRδ+) by flow cytometry of control or TRPV1-ChR2 mice optogenetically stimulated for 5 days followed by *S. mansoni* infection for 1 day. (**f**), Quantification of percentage and absolute numbers of IL-17+, (**g**), Ki67+, and (**h**), IL-13+ γδ T cells quantified by flow cytometry. *P* values were determined by two-tailed Student’s t-tests, or Two-way ANOVA with post hoc correction. *P<0.05, **P<0.01, ***P<0.001, ****P<0.0001. Representative of 2-3 independent experiments, each with ≥4 biological replicates.

### TRPV1+ neurons stimulate recruitment and activation of neutrophils and monocyte after ***S. mansoni* infection.**

Exposure to *S. mansoni* cercariae increased ear thickness in control mice, which was further exacerbated when TRPV1 neurons were stimulated prior to infection (**Fig. 3a**). Given that IL-17+ γδ T cells are well-known to drive neutrophil recruitment during inflammation^32,33^, we predicted that activation of TRPV1 neuron activation would further increase neutrophil accumulation after *S. mansoni* infection. Evaluation of myeloid cell populations by flow cytometry (**Supp. Fig. 3a**) showed that skin neutrophil and monocyte proportions and absolute numbers were increased in TRPV1-ChR2 mice compared to littermate controls 1 day following *S. mansoni* exposure (**Fig. 3b-d**). However, macrophages, Langerhan cells, type 1 and 2 conventional dendritic cells (cDC1/2) were no different between light-stimulated control or TRPV1-ChR2 mice after *S. mansoni* infection (**Supp. Fig. 3b**). Inducible nitric oxide synthase (iNOS) expression levels were analyzed in neutrophil and monocyte populations using flow cytometry. Data show that TRPV1 neuron stimulation increased the proportions and absolute numbers of iNOS+ neutrophils and monocytes in skin cell infiltrates of *S. mansoni* infected mice (**Fig. 3e,f**). In contrast, monocytes from control or TRPV1-ChR2 mice expressed similar levels of MHC-II (**Supp. Fig. 3c,d**), collectively suggesting that TRPV1 neuron activation induces neutrophil and monocyte recruitment and activation following *S. mansoni* percutaneous exposure.

**Figure 3.**
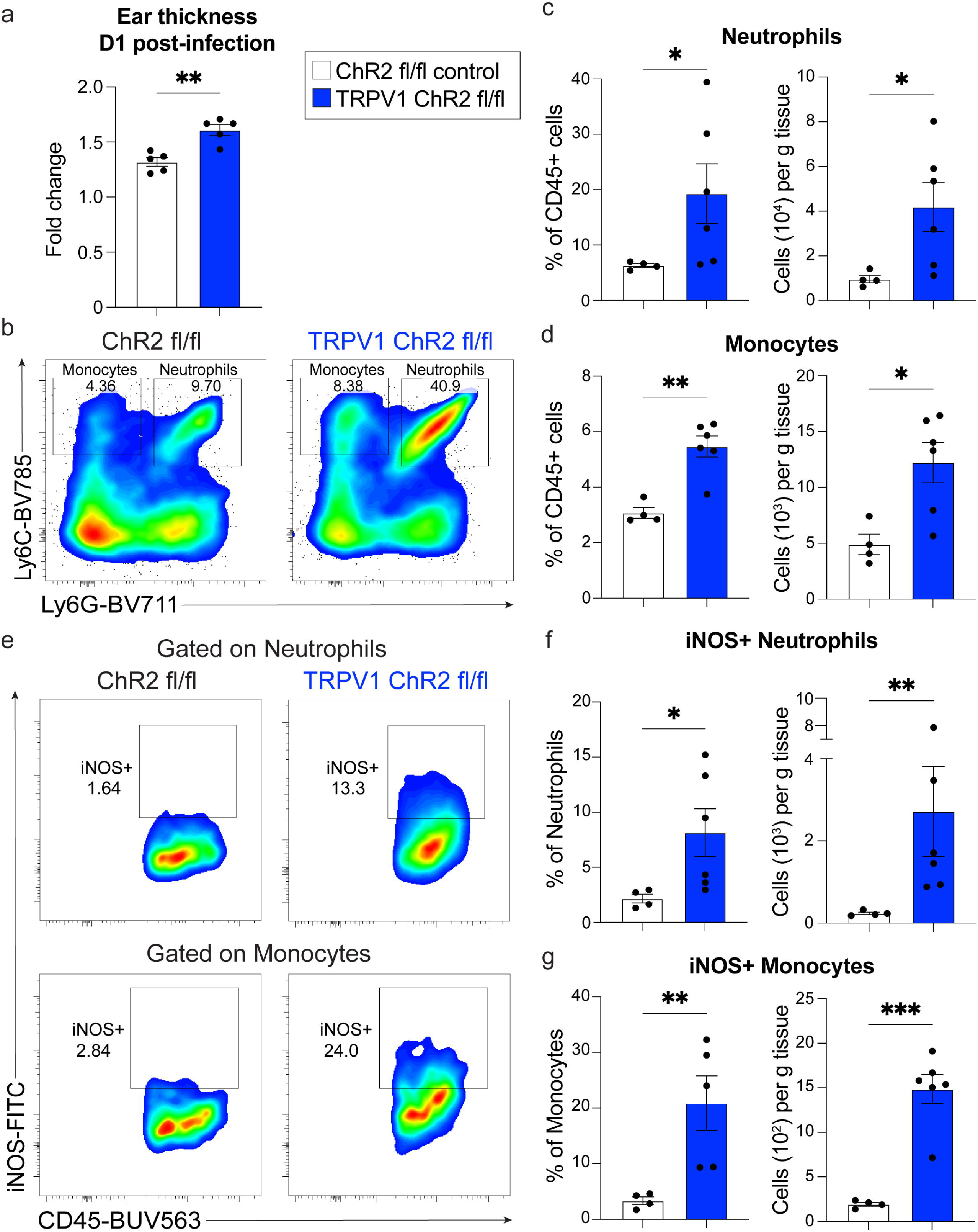
TRPV1 neurons promote neutrophil and monocyte recruitment and activation after *S. mansoni* infection. (**a**), Ear thickness of light-stimulated control or TRPV1-ChR2 mice measured 1 day after exposure to *S. mansoni* cercariae. (**b**), Representative dot plots of neutrophils (CD11b+ Ly6G+ Ly6Cint) and monocytes (CD11b+ Ly6G-Ly6Chi) assessed by flow cytometry of light-stimulated control or TRPV1-ChR2 mice 1 day after *S. mansoni* infection. (**c,d**), Percentage and absolute numbers of neutrophils and monocytes from *S. mansoni*-infected control or TRPV1-ChR2 mice. (**e-g**), Representative dot plots and quantification of iNOS expression in neutrophils and monocytes assessed by flow cytometry of light-stimulated control or TRPV1-ChR2 mice 1 day after *S. mansoni* infection. *P* values were determined by two-tailed Student’s t-test. *P<0.05, **P<0.01, ***P<0.001, ****P<0.0001. Representative of 2-3 independent experiments, each with ≥4 biological replicates.

### TRPV1 neurons are required for cutaneous host-protective responses against *S. mansoni*

To directly test whether TRPV1+ neurons contributed to host protection against *S. mansoni*, WT mice were chemically ablated of TRPV1+ afferents by subcutaneous injection with resiniferatoxin (RTX)^34^. Consistent with previous reports^1,2,5^, mice treated with RTX exhibited increased withdrawal latency to heat compared to vehicle-treated controls (**Fig. 4a**). Next, *S. mansoni*-infected mice treated with vehicle or RTX were evaluated for numbers of non-skin penetrating cercariae and lung larval numbers at 6 days post-infection. While there were no significant differences in the percentage of non-penetrating cercariae between vehicle- and RTX-treated mice, RTX treatment led to elevated lung larval burden (**Fig. 4b,c**). Moreover, RTX-treated mice had reduced IL-17+, IL-13+, and Ki67+ γδ T cells compared to vehicle-treated controls 1 day after *S. mansoni* infection (**Fig. 4d-f**). Congruent with this finding, the frequency and number of skin neutrophils and monocytes after *S. mansoni* infection were reduced in mice depleted of TRPV1+ neurons (**Fig. 4g,h**). Collectively, these results indicate that TRPV1 neurons were both essential and sufficient to promote cutaneous resistance against *S. mansoni*, primarily through inhibiting parasite dissemination to the lung.

**Figure 4.**
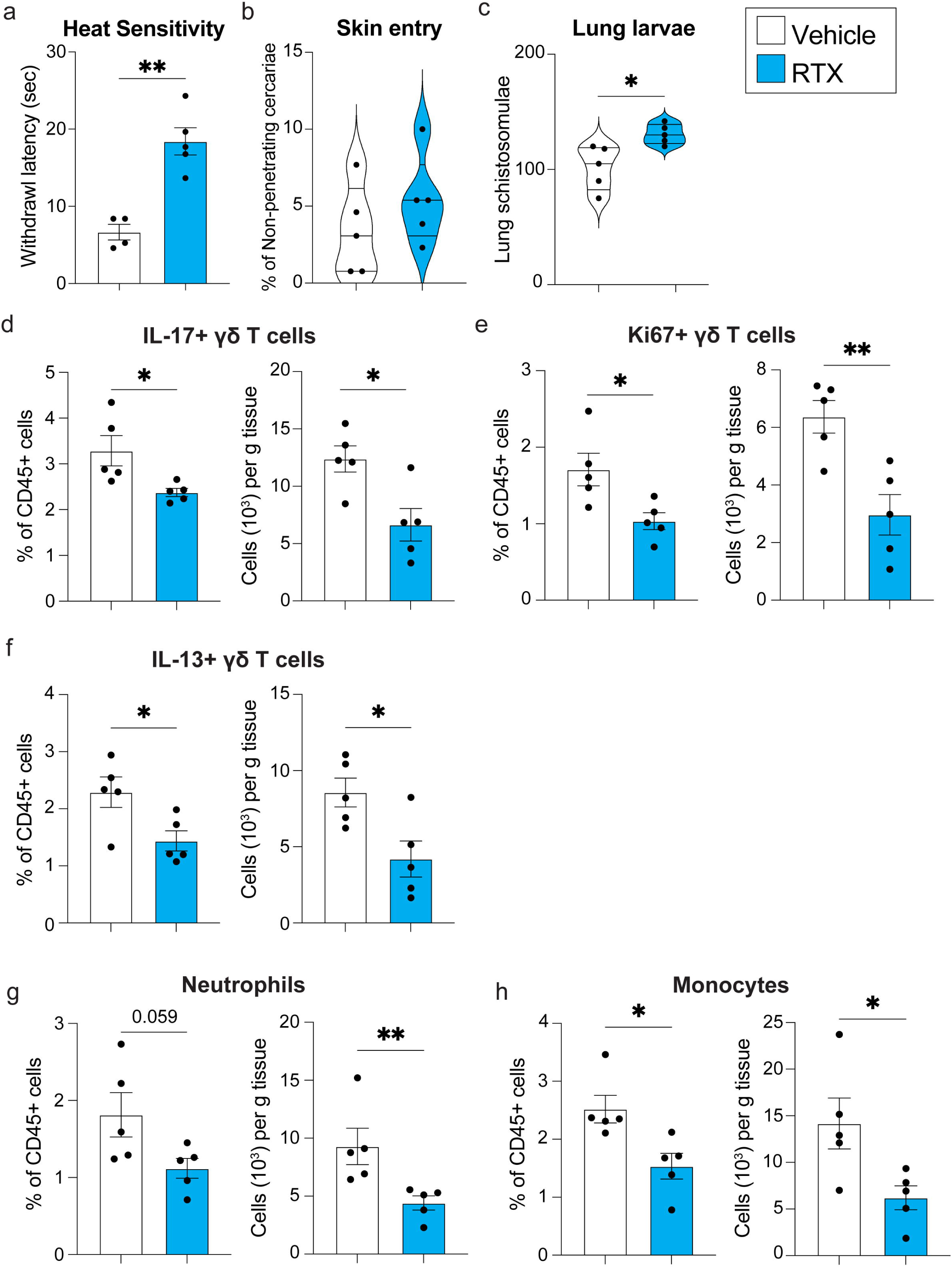
TRPV1 neurons are required to mitigate *S. mansoni* lung larval burden, γδ T cell responses and myeloid cell skin accumulation. (**a**), Thermal pain sensitivity in the paws of wildtype mice treated with vehicle or resiniferatoxin (RTX) assessed by withdrawal times (seconds) to infrared heat. (**b,c**), Percentage of non-penetrating cercariae and d6 lung larval burden from vehicle or RTX-treated mice infected with *S. mansoni.* (**d**), Percentage and absolute numbers of IL-17+, (**e**), Ki67+, and (**f**), IL-13+ γδ T cells 1 day post-infection with *S. mansoni* in vehicle or RTX-treated mice. (**g,h**), Skin neutrophils and monocytes from vehicle- or RTX-treated mice infected with *S. mansoni*. *P* values were determined by two-tailed Student’s t-test. *P<0.0 5, **P<0.01, ***P<0.001, ****P<0.0001. Representative of 2-3 independent experiments, each with ≥5 biological replicates.

## Discussion

Host-protective responses in the skin require rapid pathogen detection that elicits the recruitment and activation of hematopoietic cells, but the cellular interactions that govern this process remain undetermined. Pain caused by bacterial and fungal skin infection elicits behaviors that may promote wound healing such as aversion to touch^35^, which suggests that nociceptors may contribute to pathogen detection and clearance or even to wound healing once an infection is resolved. Curiously, *S. mansoni* cercariae seldom cause irritation or pain as they enter the skin but cause chronic human infections that last decades^16^. This study shows that pain-sensing behavior, calcium influx, and neuropeptide release evoked by TRPV1+ afferents are substantially diminished upon infection with *S. mansoni*. TRPV1 expression was not reduced in DRG neurons harvested 1 day after *S. mansoni* infection, suggesting that reduced nociceptor functions are not due to loss of TRPV1 expression. Although reduced TRPV1 neuron functions could be the result of host-derived mechanisms yet to be determined, data show that exposure to *S. mansoni* extract reduces calcium-influx^23^ and neuropeptide release induced by the TRPV1 cognate ligand, capsaicin. Upon skin encounter, *S. mansoni* cercariae quickly invade the outer layers of the epidermis and use protein secretions from their acetabular glands to degrade the dermal extracellular matrix as well as to suppress skin immunity, including serine peptidases, Sm16, and SmTeg^17,36,37^. Therefore, it is possible that *S. mansoni*-secreted molecules act on TRPV1+ afferents innervating the skin to inactivate their nociceptive functions. It may be likely that other species of parasitic helminth species that invade mammalian skin, such as hookworms and *Strongyloides* spp. have evolved related mechanisms of host modulation.

Suppression of TRPV1+ neuron activity after *S. mansoni* infection provoked the hypothesis that TRPV1+ neurons would actually promote skin immunity. Indeed, two complementary gain- and loss-of-function approaches demonstrated that TRPV1+ nociceptors promoted host-protective responses that impeded larval migration from the skin into the lungs. TRPV1 is expressed by different subsets of sensory afferents in mice, including peptidergic (PEP)1, non-peptidergic (NP)2, and NP3 neurons, as well as in keratinocytes and CD4+ T cells^38–40^. Although the effects of manipulating MrgprA3+ (NP2) neurons in the context of *S. mansoni* infection^23^ are similar to TRPV1 neuron stimulation, the relative contributions of TRPV1-expressing discrete cellular types during homeostasis and infection remains to be determined. Data also show that TRPV1 neurons were necessary and sufficient to stimulate rapid proliferation and cytokine expression by γδ T cells following *S. mansoni* infection. We and others have shown the remarkable proximity of myeloid APCs to MrgprA3+ and Nav1.8+ sensory afferents (that can also express TRPV1)^5,6,12,23^. Moreover, Nav1.8+, TRPV1+ and MrgprA3+sensory neuron activation promoted the expression of cytokines such as IL-1β, IL-23, and TNF in APCs, which have been shown to induce IL-17 secretion by lymphocyte subsets^41–43^. It is likely that TRPV1 neuron stimulation induced the γδ T cell activation through myeloid APC cytokine secretion. While earlier published studies by Mountford and colleagues concluded that skin protective responses against *S. mansoni* were required IFNγ secretion by CD4+ T cells^44,45^, whether TRPV1 neurons induce IFNγ secretion remains undetermined. We show that TRPV1 neurons promoted neutrophil and monocyte skin accumulation, possibly due to combined actions of γδ T cells and putative neuronal signals. While γδ T cell-derived IL-17 may induce keratinocytes to produce chemokines that direct neutrophil recruitment, chemokines secreted by Nav1.8+ neurons such as CCL2 may promote monocyte recruitment^6,32,33^. Unexpectedly, TRPV1 neuron stimulation induced significantly greater iNOS expression in neutrophils and monocytes. Both, neutrophils and iNOS expression are enriched in mice vaccinated against *S. mansoni* and are proposed to directly elicit killing of *Schistosoma* larvae^46–48^. We propose that TRPV1-neurons activated neutrophils and monocytes to impede parasite migration through the skin and subsequent lung dissemination. However, bacterial activation of TRPV1+ neurons causes hyperalgesia and pruritus, while inhibiting neutrophil recruitment and bacterial killing^2,8,49^, suggesting context-dependent contributions of TRPV1+ neurons to skin immunity. A better understanding of the contributions of specific sensory neuron subsets and non-neuronal cells that co-express TRPV1 (NP2 vs. NP3 vs. PEP1 vs. keratinocytes vs. CD4+ T cells) may inform effective therapies against S. mansoni infection and other cutaneous diseases. Altogether, our studies propose a model in which upon skin entry, *S. mansoni* larvae block TRPV1 neuron activation to prevent IL-17, neutrophil and monocyte responses that impede their systemic dissemination.

## MATERIALS AND METHODS

### Study design

The objectives of this study were to determine whether the mammalian infection with the trematode *Schistosoma mansoni* alters the response of TRPV1 neurons and if these pain-sensing afferents regulate cutaneous skin immunity after exposure to infectious larvae. Experiments using mice aimed for group sizes of four to eight mice (matched for age and sex) and were repeated two to three times to assure reproducibility. Sample sizes were first defined by previous experience from this and other laboratories and then corroborated by Power analysis. Mice were tattooed for identification, with experimental groups randomized across cages to account for any microisolator effects. Thermal pain sensation experiments were evaluated in 8-10 week old male mice, whereas parasite infection experiments and cell culture experiments were done using sex-matched 8-10 week old male or female mice. To address subjectivity during the study, all animal cages and experimental samples/groups were assigned a letter number code to ensure that experiments were conducted in a blinded manner. For all flow cytometry experiments, we included negative controls, such as fluorescence minus one controls, to establish reliable and reproducible gates for each marker.

### Mice and experimental procedures

C57BL/6J wildtype (WT) mice (strain #000664), B6.129-*Trpv1^tm1(cre)Bbm^*/J (TRPV1^Cre^, strain 017769)^30^, and B6.Cg-Gt(ROSA)^26Sortm32(CAG-COP4*H134R/EYFP)Hze^/J (Ai32, strain 024109) mice^31^ were purchased from the Jackson Laboratories. Optogenetic strains were bred to homozygosity and compared with age- and gender-matched controls from a separate set of breeders housed in the same facility. Co-housed Cre-negative littermate controls were used for optogenetic experiments. All mice were housed under specific pathogen-free conditions at the University of Pennsylvania. All procedures were approved by the Institutional Animal Care and Use Committee of the University of Pennsylvania (protocol 805911). Mice were euthanized by CO2 for all tissue recovery procedures following AVMA guidelines.

For chemical denervation of TRPV1 neurons, WT mice were lightly anesthetized with isoflurane (1-4%) and given three daily escalating doses of resiniferatoxin (AdipoGen Life Sciences, 30, 70, and 100μg/kg of body weight) dissolved in 1% DMSO and diluted in PBS through subcutaneous injections in the nape of the neck as reported before^1,2,5^. Control mice were treated with vehicle solution of 1% DMSO diluted in PBS on the same days. Chemical denervation was confirmed after three weeks by assessing thermal pain sensation with the Hargreaves assay as described below.

### Evaluation of thermal pain sensitivity with the Hargreaves assay

Pain sensitivity to a heat stimulus (heat hyperalgesia) was evaluated in the plantar test device (Ugo Basile)^50^. Mice were acclimated in the chambers positioned over glass pane in a quiet environment for 15-30mins at least one day before evaluating behavioral assays. An infrarred radiant heat source (intensity 32%) was placed under infected and uninfected paws and gradually increasing the temperature of the plantar surface. The threshold of pain was determined as the latency (in seconds) to evoke a response of paw withdrawal. An exposure limit of 30 s was used to prevent tissue damage. Three measurements per paw were determined for each animal measured no closer than 5 minutes, and the mean of these were reported. Pain behavior tests were performed by blinded observers who were unaware of the treatments and genotypes.

### Dorsal Root Ganglia (DRG) neuronal cultures and stimulation

Lumbar and thoracic dorsal root ganglia (DRG) from 8-10-week-old mice were isolated as previously described^51^ and then enzymatically digested at 37°C for 20mins with 2.5 mg/mL dispase II and 1.25 mg/mL collagenase A dissolved in DMEM supplemented with 1% FBS. Cells were first dissociated with a 1mL pipette tip 20-30 times and then incubated at 37°C for additional 20mins. Next, DRG were triturated with a 23G and 27G needle 3-5 times and enzymes were neutralized with F12 media supplemented with 10% FBS, 2 mM glutamine and penicillin/streptomycin. Cell suspensions were layered in a 2-phase Percoll gradient (12% over 28%) and centrifuged at 1300g for 10mins. The fraction enriched with cell bodies was lysed of red blood cells and plated in 96-well plates previously coated with 10μg/mL laminin and 20μg/mL poly-lysine dissolved in HBSS without Ca^2+/^Mg^2+^. Cells were cultured in DMEM-F12 media containing 10% FBS, 12.5mM HEPES, 1x penicillin/streptomycin, 50 ng/mL NGF (R&D Systems) and 10μM cytosine arabinoside (Sigma-Aldrich), to restrict growth of non-neuronal cells. Cultures were maintained for 2-3 days, with media changed every 2 days. NGF was removed from the media prior to stimulation.

For neuropeptide release, DRG neurons were treated with PBS (vehicle) or 3.75μg SCA for 15 minutes followed by stimulation with either media or 1μM capsaicin for 1 hour. Cell-free supernatant was collected and CGRP and Substance P concentrations were determined by EIA (Cayman chemicals, 589001, 583751) following the manufacturer’s protocol, using a Biotek plate reader to measure absorbance, and non-linear regression was used to extrapolate sample concentrations, with R-squared values of no less than 0.95.

### Calcium influx assays

DRG neuron cell cultures were obtained and cultured as described above but cultured for 2 days in 22mm coverslips previously coated with 10μg/mL laminin and 20μg/mL poly-lysine. On the day of the experiment, cells were loaded with 50 μL of 5 μM Fura-2 dye (Thermo Fisher) dissolved in 5%FBS-DMEM for 30 minutes. Then, cells were washed with Krebs-Ringer buffer (KR) and imaged with alternating 340/380nm excitation wavelengths using a Leica DMI6000 B inverted microscope and an Evolve 13 Electron-multiplying CCD camera (Teledyne Photometrics). After regions of interest for each cell were defined, baseline readings were recorded for 1 minute followed by treatment with 1μM capsaicin for 5 minutes and exposure to 70mM KCl for 1 minute. Ratiometric analysis of 340/380 signal intensities was performed only in cells where 70mM KCl induced an increase of 20% over the average signal during baseline recordings. Responsive cells were defined as described previously^12^; capsaicin-responsive neurons were considered as those with an increase of 15% from the average of baseline. For each responsive cell, Δ1340/380 ratio was normalized by dividing each 340/380 value over the 340/380 average of each cell during baseline recordings. Area under the curve (AUC) values of capsaicin-induced calcium influx were calculated with the Trapezoidal Rule. Intracellular calcium traces of Δ1340/380 Fura-2 ration of single DRG neurons were plotted using Microsoft Excel.

### Optogenetic stimulation

Mice were anesthetized with isoflurane (1-4%) and maintained on a heating pad during photostimulation. Ear thickness was assessed using calipers, taking the average of 3-5 consecutive measurements as the value for each mouse. Optogenetic stimulation was performed using a 473nm 100mW diode-pumped solid-state (DPSS) laser (SLOC Lasers, Shanghai, China), with an attached power supply and connected to a waveform generator (BX Precision, Part# 4053B), which was used to program the laser pulse conditions during stimulation. An optical cannula aimed at the dorsal ear was placed 1-2 cm from the surface of the skin and held in a fixed position with the use of a clamp. Stimulation was controlled by the waveform generator, which was programmed to deliver a sine wave pulsed at 10Hz for 30 minutes, and the experimenter adjusted the laser distance from the skin and power output to the laser to ensure a power density of 8-10 mW/mm^2^, measured periodically throughout stimulation sessions using a power meter (Thorlabs, PM100D). Mice underwent five 30-minute stimulation daily sessions.

### Isolation of *Schistosoma mansoni* cercariae

*Biomphalaria glabrata* (NMRI strain) and *Schistosoma mansoni* (NMRI strain were provided by the NIAID Schistosomiasis Resource Center of the Biomedical Research Institute (Rockville, MD) through NIH-NIAID Contract HHSN272201700014I. NIH: Biomphalaria glabrata (NMRI) exposed to Schistosoma mansoni (NMRI). Maintenance and isolation of *S. mansoni* was performed as described in^15^. Previously exposed *Biomphalaria glabrata* (NMRI strain) snails were placed under light for 1-1.5hrs at 29°C. Then supernatants containing infectious *S. mansoni* cercariae were collected, filtered, and concentrated to 100-150 cercariae in 0.1mL. For counting, live cercariae were immobilized by adding 50μL of iodine solution.

### Skin exposure to *Schistosoma mansoni* cercariae

Mice were anesthetized with Ketamine (70-100mg/kg)/Xylazine (5-12mg/kg) cocktail (i.p., 100µL/10g mouse). An infection ring was placed over the exposure area (ear pinnae) and surrounded by water-impermeable Vaseline. Then, a 100μL inoculum of 100-150 freshly isolated *S. mansoni* cercariae was placed over the skin for 20-30mins. Next, inoculum was resuspended, removed from the skin and the number of remaining cercariae present in the recovered inoculum was quantified.

### Quantification of lung *S. mansoni* parasites

For lung larvae (schistosomulae) quantification^15^, mice were infected with 500 *S. mansoni* cercariae by percutaneous infection in the ear. 6 days post-infection, lungs were recovered, mechanically digested in RPMI+10%FBS and incubated at 37°C for at least 3 hours. Then cell suspensions were passed through a cheesecloth. Larval burden was determined by microscopic quantification.

### Generation of *S. mansoni* cercarial antigens

5-10 000 *S. mansoni* cercariae were collected as described above and placed on ice for 30 minutes followed by centrifugation at 2000rpm for 5 minutes at 4C. After water was removed, parasites were washed 3 times with PBS containing 1x penicillin/streptomycin, and parasites were resuspended in PBS containing peptidase inhibitors (vehicle) and homogenized via sonication (3X, 30s, 30W). Cercarial homogenates were centrifuged at 16 000g and the soluble fraction was collected and filtered through 0.22μm membranes followed by LPS removal using Pierce High-Capacity Endotoxin Removal Resin kit (ThermoFisher, 88275) following the manufacturer’s protocol. Protein concentration was determined by Pierce BCA Protein Assay (ThermoFisher, 23225). Cercarial antigens were stored at −80C until use.

### Skin digestion and preparation for flow cytometric analysis

Ear skin explants were surgically dissected and processed similar to previously reported. Skin biopsies were mechanically dissociated and then incubated with skin digest solution containing 2mg/ml of collagenase XI (Sigma-Aldrich), 0.5mg/ml hyaluronidase (Sigma-Aldrich), and 0.1 mg/ml DNase (Roche) dissolved in DMEM-F12 containing 2.5% FBS for 25mins at 37°C with constant agitation. Skin homogenates were then passed through an 18-gauge needle 3-5 times and incubated at 37°C for additional 20mins. Then, cell preparations were passed again through an 18-gauge needle and filtered twice through 100μm and 40μm filters. Cells were then resuspended in RPMI supplemented with 10% FBS and incubated with Cell Activation Cocktail (with Brefeldin A) for 5-6hours for measurement of intracellular cytokine expression. Cell suspensions were stained for live/dead cell exclusion using LIVE/DEAD™ Fixable Aqua Dead Cell Stain Kit according to manufacturer’s protocol followed by Fc Block for 15mins at 4C and surface marker staining for 25 mins at 4C. Next, eBioscience™ Foxp3 / Transcription Factor Staining Buffer Set was used according to manufacturer’s protocol. Antibodies for nuclear proteins (Foxp3, GATA3) were incubated overnight at 4C. Intracellular staining was done for 1-1.5hrs on ice. Cells were analyzed with a BD Symphony A3 Lite.

### Immunofluorescence

DRG neurons were seeded in tissue chambers and fixed in cold 4% PFA for 2 hrs at room temperature, washed with 1XPBS for 1-4 hrs, and left in PBS containing 30% sucrose overnight. Cells were first incubated for 90 minutes at room temperature in blocking buffer (1XTBS containing 5% normal donkey serum, Fc Block, 1%BSA, and 0.3% Triton X-100). After rinsing three times with 1XTBS, sections were incubated in primary antibodies diluted in blocking buffer at 4°C overnight. Sections were rinsed and incubated with secondary antibodies diluted in blocking buffer for 120 min. at room temperature. Sections were incubated with DAPI diluted in 1XTBS 1%BSA for 10-15 min. at room temperature. Sections were mounted with Fluoroshield mounting medium. The primary antibodies used for immunofluorescence staining are guinea pig anti-mouse TRPV1 Polyclonal Antibody (1:500; ThermoFisher PA1-29770); rabbit anti-mouse sodium channel pan antibody (1:500; ThermoFisher PA5-77723). The secondary antibodies used are Alexa Fluor 488 Donkey anti-guinea pig, and Cy3 Donkey anti-rabbit at 1:500 dilution. Detection of fluorescence was observed under DAPI, GFP, Cy3 filters on a Leica DM6000 motorized upright microscope. Exposure times and fluorescence intensities were normalized to no-primary antibody control images. We photographed fluorescence channels separately, merged them together, and overlaid them atop the corresponding images.

### Statistics

Results are shown as the mean ± s.e.m. P < 0.05 was considered as significantly different. Grubb’s test was used to identify outliers and none were found. Normal distribution of the data was determined by Shapiro-Wilk test. When these data were normally distributed, we used parametric statistical tests; otherwise non-parametric analysis was used when stated. Statistical analysis was performed using Student’s t-test for two groups, one-way ANOVA for three groups, or two-way ANOVA (ear thickness measurements), with appropriate post-hoc tests in Prism 10 (GraphPad Software).

| RESOURCES TABLE |  |  |
| --- | --- | --- |
| REAGENT OR RESOURCE | SOURCE | IDENTIFIER |
| **Antibodies** |  |  |
| BUV563<sup>TM</sup> Rat Anti-Mouse CD45 Antibody | BD Biosciences | Cat# 752412. RRID:AB_2873124 |
| BUV737<sup>TM</sup> Rat Anti-Mouse CD11b Antibody | BD Biosciences | Cat# 612801. RRID:AB_2738811 |
| Brilliant Violet 605<sup>TM</sup> anti-Mouse CD11c Antibody | Biolegend | Cat# 117334. RRID: AB_2562415 |
| Brilliant Violet 650<sup>TM</sup> anti-Mouse XCR1 Antibody | Biolegend | Cat# 148220. RRID: AB_2566410 |
| Brilliant Violet 711<sup>TM</sup> anti-Mouse Ly6G Antibody | Biolegend | Cat# 127643. RRID: AB_2565971 |
| Brilliant Violet 785<sup>TM</sup> anti-Mouse Ly6C Antibody | Biolegend | Cat# 128041. RRID: AB_2565852 |
| Alexa Fluor® 700 anti-mouse I-A/I-E Antibody | Biolegend | Cat# 107622. RRID: AB_493727 |
| APC/Cy7 anti-Mouse CD326 (Ep-CAM) Antibody | Biolegend | Cat# 118218. RRID: AB_2098648 |
| PE/Dazzle<sup>TM</sup>594 anti-mouse CD172a (SIRPα) Antibody | Biolegend | Cat# 144016. RRID: AB_2565280 |
| PE/Cyanine5 anti-mouse F4/80 Antibody | Biolegend | Cat# 123112. RRID: AB_893482 |
| PE/Cyanine7 anti-mouse CD64 (FcγRI) Antibody | Biolegend | Cat# 139314. RRID: AB_2563904 |
| BUV395<sup>TM</sup> Rat Anti-Mouse IL-17 Antibody | BD Biosciences | Cat# 565246. RRID:AB_2722575 |
| BUV737<sup>TM</sup> Rat Anti-Mouse CD4 Antibody | BD Biosciences | Cat# 612844. RRID:AB_2738811 |
| Brilliant Violet 421<sup>TM</sup> anti-mouse IL-33Rα (IL1RL1, ST2) Antibody | Biolegend | Cat# 145309. RRID: AB_2565634 |
| Brilliant Violet 605<sup>TM</sup> anti-mouse TCR $\gamma/\delta$ Antibody | Biolegend | Cat# 118129. RRID: AB_2563356 |
| Brilliant Violet 711<sup>TM</sup> anti-Mouse CD90.2 Antibody | Biolegend | Cat# 105349. RRID: AB_2800564 |
| Brilliant Violet 785<sup>TM</sup> anti-mouse IL-33Rα (IL1RL1, ST2) Antibody | Biolegend | Cat# 145321. RRID: AB_2860702 |
| Gata-3 Monoclonal Antibody (TWAJ), Alexa Fluor<sup>TM</sup> 488, eBioscience<sup>TM</sup> | ThermoFisher | Cat# 53-9966-42. RRID: AB_2574493 |
| PerCP/Cy5.5 anti-Mouse CD19 Antibody | Biolegend | Cat# 152406. RRID: AB_2629815 |
| APC anti-mouse IFN-γ Antibody | Biolegend | Cat# 505810. RRID: AB_315404 |
| Ki-67 Monoclonal Antibody (SolA15), APC-eFluor<sup>TM</sup> 780, eBioscience<sup>TM</sup> | ThermoFisher | Cat# 47-5698-82. RRID: AB_2688065 |
| FOXP3 Monoclonal Antibody (FJK-16s), PE, eBioscience™ | ThermoFisher | Cat#12-5773-82.RRID:AB_465936 |
| PE/Dazzle™ 594 anti-mouse CD8a Antibody | Biolegend | Cat# 100762. RRID: AB_22564027 |
| PE/Cyanine5 anti-mouse TCR β chain Antibody | Biolegend | Cat# 109210. RRID: AB_313433 |
| IL-13 Monoclonal Antibody (eBio13A), PE-Cyanine7, eBioscience™ | ThermoFisher | Cat# 25-7133-82. RRID: AB_2573530 |
| TRPV1 Polyclonal Antibody | ThermoFisher | Cat# PA1-29770 |
| Pan sodium channel polyclonal Antibody | ThermoFisher | Cat# PA5-77723 |
| Purified anti-Loricrin Antibody | Biolegend | Cat# 905104. RRID: AB_2616895 |
| Anti-Desmoglein 1/DSG1 antibody [EPR6766(B)] | Abcam | Cat# ab124798 |
| Anti-Ki67 antibody | Abcam | Cat# ab15580 |
| DAPI Solution | ThermoFisher | Cat# 62248 |
| Cy3 AffiniPure F(ab')<sub>2</sub> Fragment Donkey Anti-Rabbit IgG (H+L) | Jackson ImmunoResearch Laboratories Inc. | Cat# 711-166-152 |
| Alexa Fluor 488 AffiniPure F(ab')<sub>2</sub> Fragment Donkey Anti-Guinea Pig IgG (H+L) | Jackson ImmunoResearch Laboratories Inc. | Cat# 703-545-155 |
| <u>Chemicals, peptides, and recombinant proteins</u> |  |  |
| Resiniferatoxin | AdipoGen | Cat# AG-CN2-0534-MC01 |
| Collagenase from *Clostridium histolyticum* | Sigma-Aldrich | Cat# C7657 |
| Hyaluronidase from bovine testes | Sigma-Aldrich | Cat# H3506 |
| DNase I from bovine pancreas | Sigma-Aldrich | Cat# 11284932001 |
| Cell Activation Cocktail (with Brefeldin A) | Biolegend | Cat# 423304 |
| Collagenase A from *Clostridium histolyticum* | Sigma-Aldrich | Cat# 10103578001 |
| Collagenase D | Roche | Cat# 11088866001 |
| Pronase from *Streptomyces griseus* | Sigma-Aldrich | Cat# 10165921001 |
| Dispase II | Sigma-Aldrich | Cat# D4693 |
| Laminin mouse protein | ThermoFisher | Cat# 23017015 |
| Poly-D-Lysine | ThermoFisher | Cat# A3890401 |
| BD GolgiPlug™ Protein Transport Inhibitor (containing Brefeldin A) | BD Biosciences | Cat# 555029 |
| BD GolgiStop™ Protein Transport Inhibitor (containing Monensin) | BD Biosciences | Cat# 554724 |
| Capsaicin | Sigma-Aldrich | Cat# M2028 |
| eBioscience™ Foxp3 / Transcription Factor Staining Buffer Set | ThermoFisher | Cat# 00-5523-00 |
| DMEM/F-12, powder | ThermoFisher | Cat# 12500062 |
| RPMI 1640 Medium, powder | ThermoFisher | Cat# 31800-022 |
| Percoll | Sigma-Aldrich | Cat# P1644 |
| Trypsin-EDTA (0.05%), phenol red | ThermoFisher | Cat# 25300054 |
| Maxima H Minus Reverse Transcriptase | ThermoFisher | Cat# EP0753 |
| Recombinant Mouse beta-NGF Protein | R&D systems | Cat# 1156-NG-100/CF |
| Cytosine β-D-arabinofuranoside hydrochloride | Sigma-Aldrich | Cat# C6645 |
| 16% Paraformaldehyde (formaldehyde) aqueous solution | Electron Microscopy Sciences | Cat#15710-S |
| Normal donkey serum | ThermoFisher | Cat# NC9624464 |
| Fluoroshield Histology Mounting Medium | Sigma-Aldrich | Cat# F6182 |
| Ketaset (Ketamine hydrochloride injection) | Zoetis | NADA# 043-304 |
| Anased (Xylazine injection) | AKORN | NADA# 139-236 |
| Fura-2 AM, cell permeant | ThermoFisher | Cat# F1201 |
| <u>Commercial Assays</u> |  |  |
| Substance P ELISA Kit | Cayman Chemicals | Cat# 583751 |
| CGRP (rat) EIA Kit | Cayman Chemicals | Cat# 589001 |
| LIVE/DEAD™ Fixable Aqua Dead Cell Stain Kit, for 405 nm excitation | ThermoFisher | Cat# L34966 |
| Pierce High-Capacity Endotoxin Removal Resin kit | ThermoFisher | Cat# 88275 |
| Pierce BCA Protein Assay kit | ThermoFisher | Cat# 23225 |
| <u>Experimental models: Organisms/strains</u> |  |  |
| Mouse: C56BL/6J | The Jackson Laboratory | Strain # 000664 |
| Mouse: B6.129-*Trpv1*<sup>tm1(cre)Bbm</sup>/J | The Jackson Laboratory | Strain #017769 |
| Mouse: B6.Cg-Gt(ROSA)<sup>26Sortm32</sup>(CAG-COP4*H134R/EYFP)<sup>Hze</sup>/J (Ai32) | The Jackson Laboratory | Strain #024109 |
| Parasite: *Schistosoma mansoni* | NIAID Schistosomiasis Resource Center | Strain: NMRI |
| Snails: *Biomphalaria glabrata* | NIAID Schistosomiasis Resource Center | Strain: NMRI |
| <u>Instruments and equipment</u> |  |  |
| Plantar test (Hargreaves method) | Ugo Basile | Sku:37570 |
| Symphony A3 Lite | BD Biosciences | N/A |
| Aria II | BD Biosciences | N/A |
| MITUTOYO Dial Thickness Gauge | Grainger | Part # 6NRC7 |
| 473nm DPSS Laser Nd:YAG | SLOC lasers | Model BL473T8-100FC |
| Power source | SLOC lasers | Model ADR-800A |
| 1x2 MM Fiber Optic Coupler | Thor Labs | Part# TM200FS1B |
| Power and energy meter | Thor Labs | Part# PM100D |
| 10 MHz Dual Channel Function/Arbitrary Waveform Generator | BX Precision | Part# 4053B |
| Leica DM6000 | Leica | N/A |
| Leica DMI6000 B | Leica | N/A |
| Evolve 13 Electron-multiplying CCD camera | Teledyne Photometrics | N/A |
| Leica TCS SP8 WLL Confocal | Leica | N/A |
| <u>Software and algorithms</u> |  |  |
| Flowjo 10.8.1 | Flowjo | N/A |
| Prism 10 | Graph Pad | N/A |
| Leica Application Suite X Version 5.1.0.25446 | Leica | N/A |

## Acknowledgments

We thank William Wilkerson and the TRIP summer research program for supporting his contribution to this work. We thank the NIAID Schistosomiasis Resource Center of the Biomedical Research Institute (Rockville, MD) for providing Biomphalaria glabrata (NMRI) exposed to *Schistosoma mansoni* (NMRI) through NIH-NIAID Contract HHSN272201700014I. This work was supported by the National Institutes of Health (grant nos. T32 AI007532-24, RO1 AI164715-01, U01 AI163062-01, RO1 AI123173-05 to D.R.H). J.M.I.R. is supported by the Life Sciences Research Foundation and by the Skin Biology and Disease Research Core Pilot and Feasibility Grant of the University of Pennsylvania.

## Author contributions

Conceptualization: JMIR, DRH

Methodology: JMIR, CN, LYH, CFP, AS, HLR

Investigation: JMIR, CN, LYH, CFP, AS, HLR

Visualization: JMIR, AS, DR

Funding acquisition: JMIR, DRH

Supervision: WL, DRH

Writing – original draft: JMIR, AS

Writing – review & editing: JMIR, AS, DRH

## Lead contact

Further information and requests for reagents may be directed to, and will be fulfilled by, the lead contact De’Broski R. Herbert.

## Data, Code, and Materials availability

All reagents generated or used in this study are available on request from the lead contact with a completed Materials Transfer Agreement. Information on reagents used in this study is available in the key resources table.

---

**Supplementary Figure 1.**
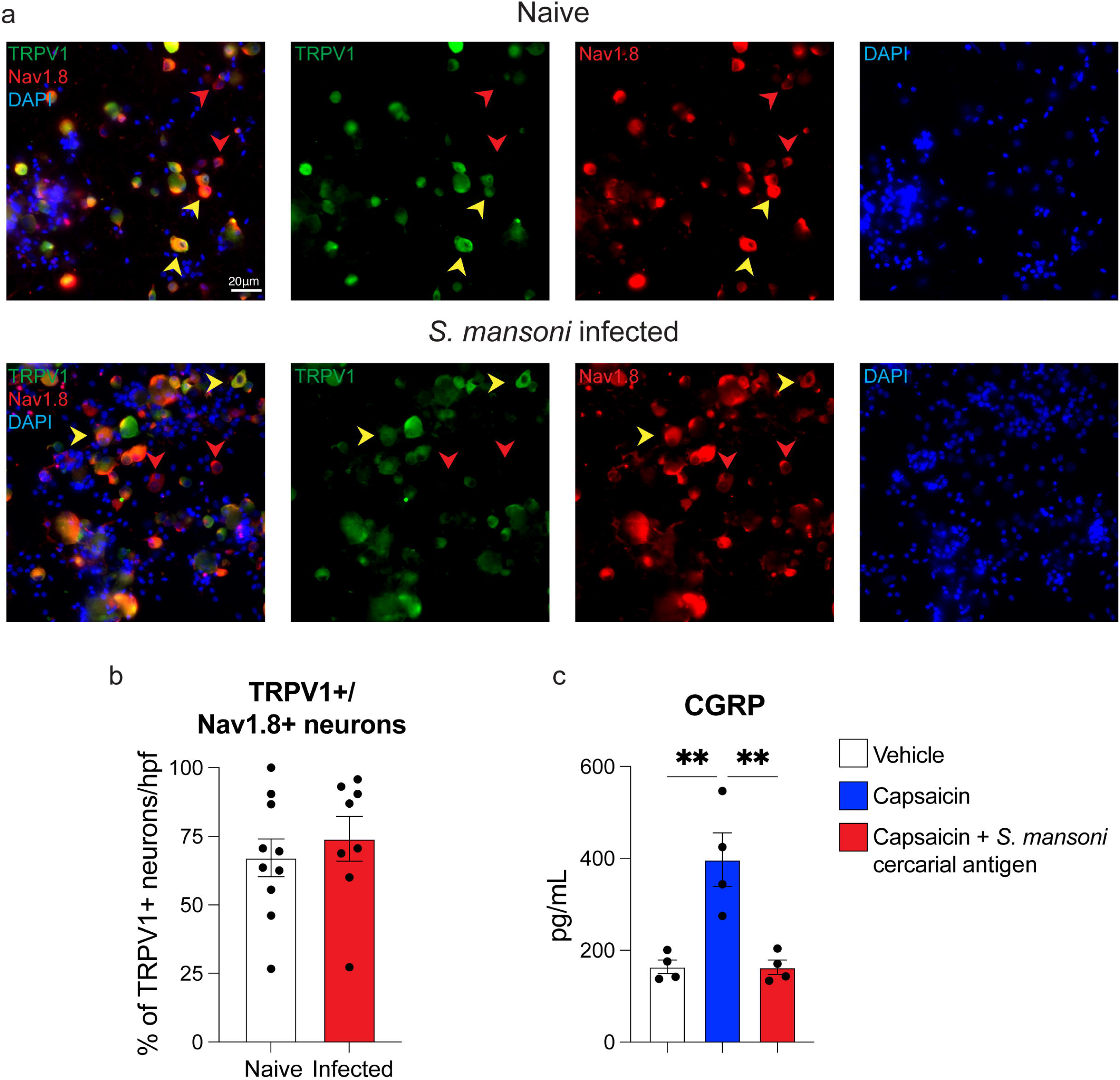
*S. mansoni* infection does not alter TRPV1 expression in DRG neurons. (**a,b**), Immunofluorescence (IFA) staining and quantification of TRPV1 (green) and Nav1.8 (red) expression in DRG neurons from naïve or *S. mansoni* infected mice. Yellow arrows indicate TRPV1 and Nav1.8 co-expression. Red arrows indicate Nav1.8 expression. (**b**), CGRP levels from DRG neurons treated with vehicle or *S. mansoni* cercarial extract for 15 minutes prior to capsaicin stimulation for 1 hour. *P* values were determined by two-tailed Student’s t-test or One-way ANOVA with post hoc correction. **P<0.01. Representative of 2-3 independent experiments, each with ≥4 biological replicates.

**Supplementary Figure 2.**
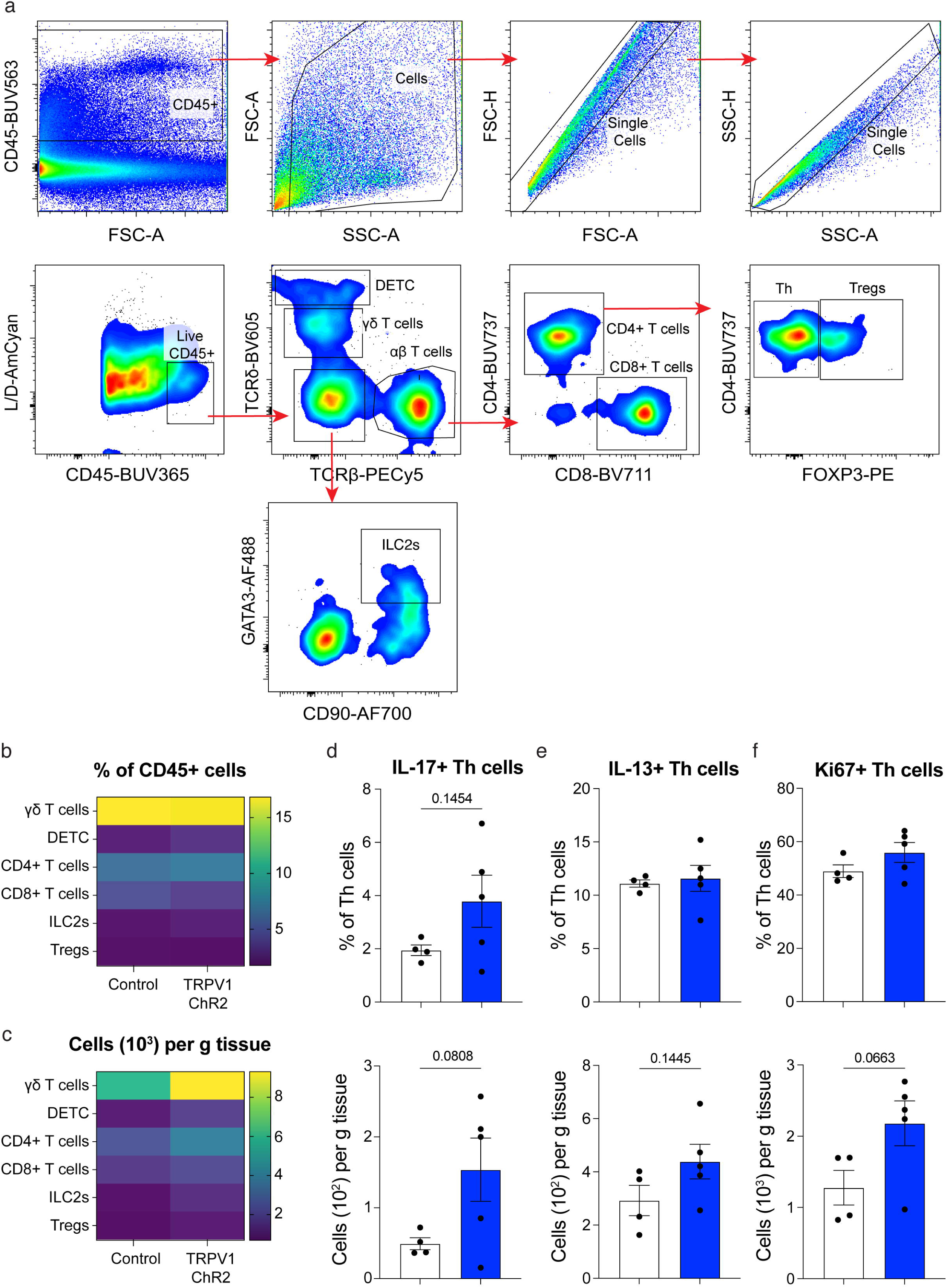
Evaluation of lymphoid cell populations in control or TRPV1-ChR2 mice after *S. mansoni* infection. (**a**), Gating strategy to analyze skin-resident lymphoid cell populations by flow cytometry. (**b,c**), Heatmaps representing the average of percentage and absolute numbers of skin γδ T cells, dendritic epidermal T cells (DETC), CD4+ T helper cells, CD8+ T cytotoxic cells, group 2 innate lymphoid cells (ILC2s) and T regulatory cells (Tregs) in control or TRPV1-ChR2 mice 1 day post *S. mansoni* infection. (**d**), Percentage and absolute numbers of IL-17+, (**e**), Ki67+, and (**f**), IL-13+ CD4+T helper cells 1 day post-infection with *S. mansoni* in control or TRPV1-ChR2 mice. *P* values were determined by two-tailed Student’s t-test. Representative of 2-3 independent experiments, each with ≥4 biological replicates.

**Supplementary Figure 3.**
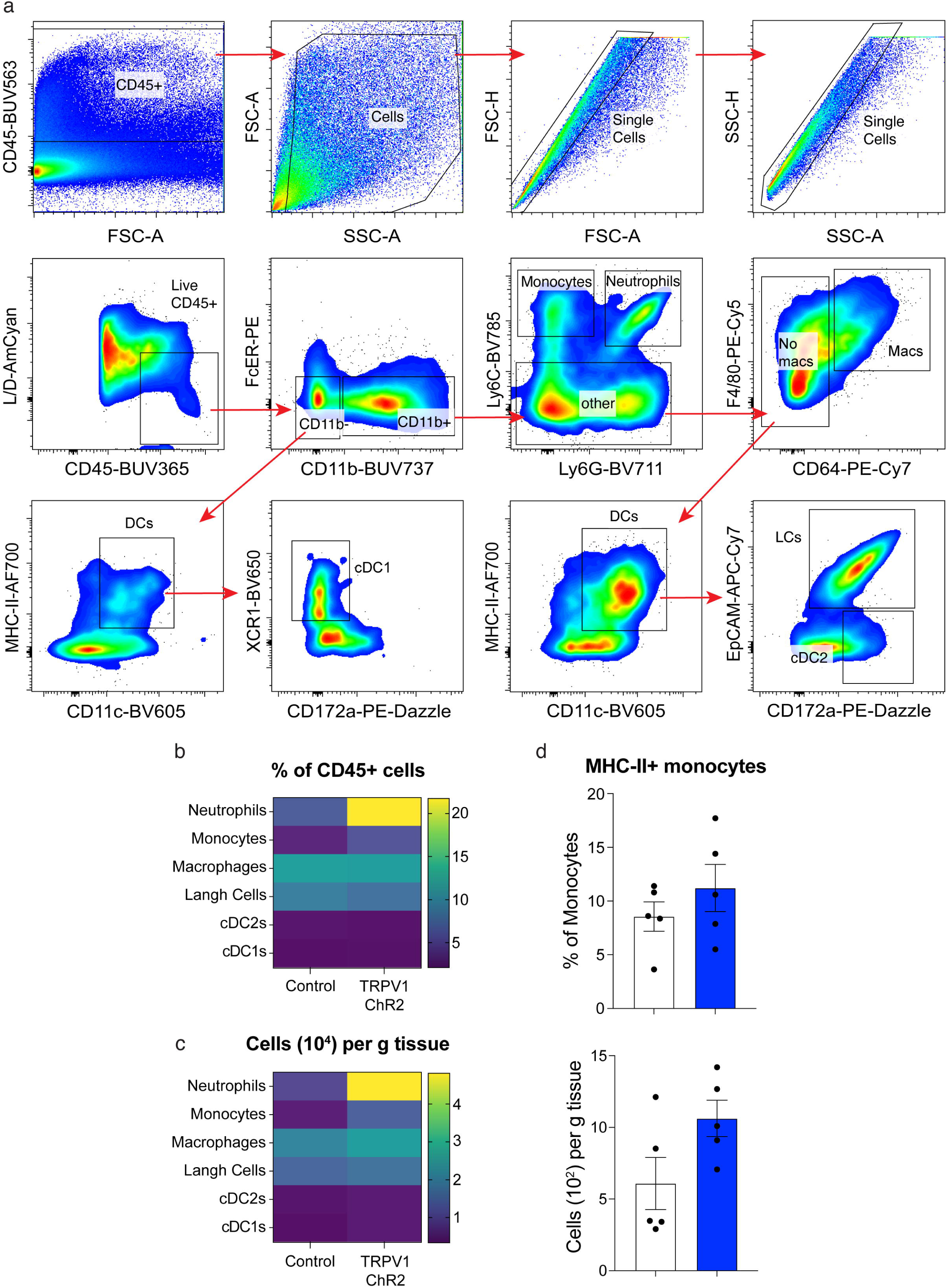
Evaluation of myeloid cell populations in control or TRPV1-ChR2 mice after *S. mansoni* infection. (**a**), Gating strategy to analyze skin-resident myeloid cell populations by flow cytometry. (**b,c**), Heatmaps representing the average of percentage and absolute numbers of skin neutrophils, monocytes, macrophages, Langerhan Cells, type 1 conventional dendritic cells (cDC1), and cDC2s in control or TRPV1-ChR2 mice 1 day post *S. mansoni* infection. (**d**), Percentage and absolute numbers of MHC-II expression in monocytes 1 day post-infection with *S. mansoni* in control or TRPV1-ChR2 mice. *P* values were determined by two-tailed Student’s t-test. **P<0.01. Representative of 2-3 independent experiments, each with ≥4 biological replicates.

